# Low Dose Lead Exposure Induces Alterations on Heterochromatin Hallmarks Persisting Through SH-SY5Y Cell Differentiation

**DOI:** 10.1101/2020.07.27.224246

**Authors:** Li Lin, Junkai Xie, Oscar F. Sanchez, Chris Bryan, Jennifer Freeman, Chongli Yuan

## Abstract

Lead (Pb) is a commonly found heavy metal due to its historical applications. Recent studies have associated the early-life Pb exposure with the onset of various neurodegenerative disease. The molecular mechanisms of Pb conferring long-term neurotoxicity, however, is yet to be elucidated. In this study, we explored the persistency of alteration in epigenetic marks that arise from exposure to low dose of Pb using a combination of image-based and gene expression analysis. Using SH-SY5Y as a model cell line, we observed significant alterations in global 5-methycytosine (5mC) and histone 3 lysine 27 tri-methylation (H3K27me3) and histone 3 lysine 9 tri-methylation (H3K9me3) levels in a dose-dependent manner immediately after Pb exposure. The changes are partially associated with alterations in epigenetic enzyme expression levels. Long term culturing (14 days) after cease of exposure revealed persistent changes in 5mC, partial recovery in H3K9me3 and overcompensation in H3K27me3 levels. The observed alterations in H3K9me3 and H3K27me3 are reversed after neuronal differentiation, while reduction in 5mC levels are amplified with significant changes in patterns as identified via texture clustering analysis. Moreover, correlation analysis demonstrates a strong positive correlation between trends of 5mC alteration after differentiation and neuronal morphology. Collectively, our results suggest that exposure to low dose of Pb prior to differentiation can result in persistent epigenome alterations that can potentially be responsible for observed phenotypic changes.

## 1. INTRODUCTION

Lead (Pb) is a heavy metal historically used for various applications, i.e., painting, ceramic glazing, pipe soldering, and gasolines (Nriagu, 1990; Wilson and Horrocks, 2008) and is thus found in soil and water sources due to its historical uses (Annest et al., 1983). A survey conducted in late 1970s suggested that ~ 78% of the population then has blood lead levels (BLLs) of 10 μg/dL (100 parts per billion, ppb) or higher; and this percentage is even higher in children (1-5 years old) with an estimated percentage of 88% (Dignam et al., 2019). Usage of Pb has thus been tightly regulated by environmental protection agencies. High Pb levels exceeding the regulatory standard, however, are often reported in many geographic locations due to legacy uses. Human exposure to Pb can primarily be attributed to ingestion of contaminated food (~ 59 – 81% of total Pb exposure cases(Watanabe et al., 2000)) or water (~20% of total (Jarvis et al., 2018)).

Pb exposure can happen at different life stages. Among them, developmental exposure to Pb is often associated with more detrimental health outcomes, including severe damages in central nervous system (Nevin, 2000, 2007) and thus has attracted significant attention in recent years. Specifically, children with prenatal Pb exposure (> 100 ppb) exhibited delayed cognitive development (Bellinger et al., 1987), while children exposed to Pb at young ages show severe impairment in their cognitive functions and often poor academic performances(Cecil et al., 2008b). Interestingly, post-natal exposure to equivalent concentrations of Pb seems to cause more significant damages than prenatal exposure (Bellinger et al., 1987; Bellinger et al., 1992; Lanphear et al., 2000). Pathologically, exposure to Pb during childhood (< 6.5 years old) can cause significant changes in brains at adulthood (19 – 24 years old) including alterations in brain volume, architecture and metabolism(Cecil et al., 2008a) (Cecil et al., 2011). Recent studies have further associated early-life Pb exposure to various neurodegenerative disorders, such as Alzheimer’s disease (AD) (Eid et al., 2016; Bihaqi et al., 2017) and Parkinson’s disease (PD) (Weisskopf et al., 2010; Caudle et al., 2012) aligning with the Developmental Origins of Health and Disease (DOHaD) paradigm. Collectively, these results have led to the recent revision of action doses of Pb in drinking water from 50 to 15 ppb by the U.S. Environmental Protection Agencies.

The neurotoxicity of Pb, in the form of Pb^2+^, partially arises from its resemblance to other cations essential for neuron activities, such as Ca (Ca^2+^) and Na (Na^+^) (Needleman, 2004). Specifically, Pb competes with Ca in neuronal signaling pathways (Flora et al., 2012; Meissner, 2017) and can disrupt the excitatory and inhibitory synaptic transmission balance in hippocampal neurons(Zou et al., 2020). Pb can also interfere with Na uptake and subsequently affect neuron firing(Yan et al., 2008). The long-term health implications of Pb exposure, particularly increased risk for AD and PD (Coon et al., 2006; Mansouri and Cauli, 2009; Weisskopf et al., 2010; Caudle et al., 2012; Eid et al., 2016; Bihaqi et al., 2017), however, cannot be fully explained by such a mechanism. Moreover, Pb exposure is shown to have transgenerational effects in altering brain transcriptome by a recent zebrafish study(Meyer et al., 2020) suggesting the involvement of epigenetic mechanism accounting for low mutagenic activities of Pb at low doses.

Recent studies have been associating exposure to Pb with epigenome changes. For example, developmental Pb exposure in mice can cause expression changes in DNA methyltransferase 1 (DNMT1), DNA CpG methylation (^me^CpG) and other histone modification levels, including H3K9ac, H3K4me2 and H3K27me3(Eid et al., 2016). Similar observations were also made in rats (Singh et al., 2018a). Pb can also affect the activities of epigenetic enzymes, i.e., DNMTs as shown in our previous study (Sanchez et al., 2017). Few studies exist, however, assessing the persistence of Pb-induced epigenetic changes over time. It thus remains elusive as to how epigenetic changes induced by Pb exposure can persist through the developmental stage and confer long-term health risks after the cessation of exposure to Pb.

Here, we worked with SH-SY5Y cell line, a model human neuroblastoma cell line capable of differentiating into neurons (Xicoy et al., 2017) to examine epigenetic changes induced by exposure to Pb that potentially persist after the cessation of exposure. Exposure to low doses of Pb (15 and 50 ppb) prior to differentiation is found to induce significant changes in the percentage of mature neurons as well as neurite lengths. Significant epigenetic changes were observed right after Pb exposure potentially attributing to expression changes in epigenetic enzyme. Changes in 5mC levels persist after cessation of exposure and amplifies through neuronal differentiation, suggesting a potential correlation with the observed nonlinear dose-dependence of phenotypical alteration.

## 2. MATERIAL AND METHODS

### 2.1 Culture and differentiation of SH-SY5Y

SH-SY5Y (ATCC, CRL-2266) was cultured following the standard protocol recommended by the manufacturer. Specifically, cells were maintained in Eagle’s Minimum Essential Medium (Gibco, U.S.) supplemented with 10% (v/v) fetal bovine serum (Atlanta Bio, U.S.), 1% of Penicillin-Streptomycin and 1% of MEM Non-Essential Amino Acids (Gibco, U.S.). Cells were passed in T25 flasks at 37 °C with 5% CO_2_.

SH-SY5Y differentiation was carried out following a protocol adapted from literature (Shipley et al., 2016) (see also **Fig. 1A)**. Briefly, differentiation was started on Day 0 by an exchange of culture medium, proceeded to Day 6 with a second medium exchange and completed on Day 14. Compositions of differentiation medium can be found in **Table S1** (Supplementary information). The completion of differentiation was confirmed by staining cells with MAP2-antibody (PA5-17646, Invitrogen, U.S.) that are specific for mature neurons.

**Fig. 1.**
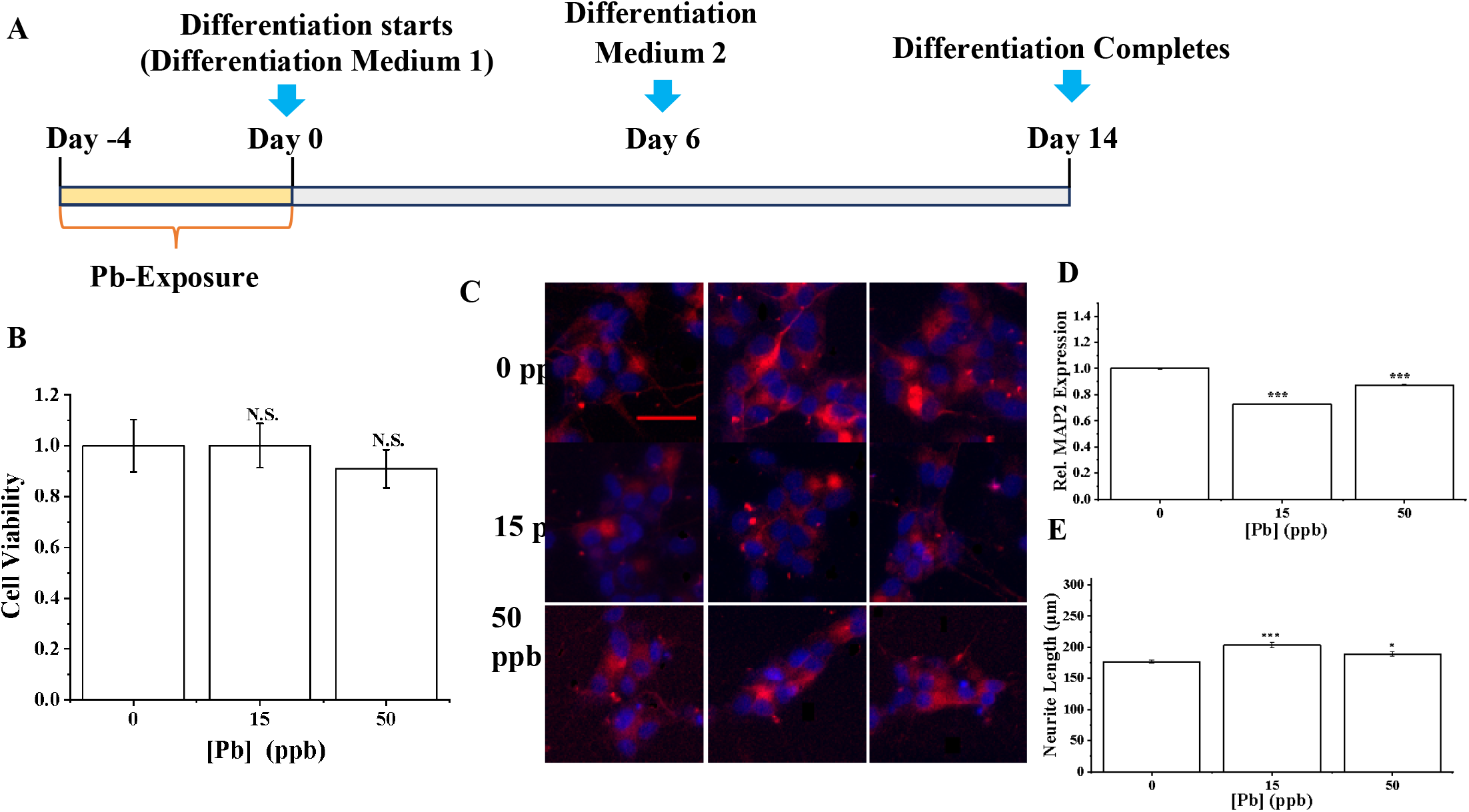
**(A)** A schematic illustration of the exposure and differentiation timeline for SH-SY5Y cells. **(B)** Viability of cells after exposure to Pb of different doses for 96 hours assayed using MTT. Data = mean ± standard deviation. n = 3. (C)Differentiated SH-SY5Y cells (Day 14) were stained with MAP2 antibody (red) and DAPI (blue). Cells were exposed to various concentrations of Pb prior to the initiation of differentiation. Scale bar = 50 μm **(D)** A bar plot showing relative MAP2 expression levels in Pb-naïve and treated cells. **(E)** Neurite length of differentiated SH-SY5Y neurons that have been exposed to Pb of varying doses. In **(D)** and **(E)**, n > 300 cells. N.S.: no significance (p > 0.05). * [=] p < 0.05, *** [=] p < 0.001. Data = mean ± standard error.

### 2.2 Selection of Pb concentrations and treatment

Pb stock solutions were prepared as described previously(Lee et al., 2017) and spiked into cell culture medium at selected concentrations. We have chosen Pb concentrations of 15 and 50 ppb in this work based on the past and current EPA regulation standard on drinking water (50 and 15 ppb, respectively)(2007). Pb concentrations higher than 15 ppb have also been reported in recent public health incidence (e.g., 16.5-33.2 μg L^−1^ (ppb) (Pieper et al., 2018)).

All cells were treated with Pb of 0, 15 and 50 ppb for 96 hours prior to differentiation (see also **Fig. 1A**) or continuous culturing. All cells were washed three times in PBS prior to the initiation of differentiation or continuous culturing.

### 2.3 Cell viability and morphological assessments

Cell viability was assessed using a colorimetric MTT assay (ab211091, Abcam, U.S.). Nuclear size and morphology were determined by first staining cell nucleus with Draq5 (Thermo Fisher, U.S.) followed by imaging using fluorescent microscopy (Nikon, Japan). Individual nucleus was identified using a customized CellProfiler (Broad Institute, U.S.) pipeline as we detailed in our previous work(Sánchez et al., 2020).

Neurite length was determined via cell images collected using a bright-field microscope (Floid, Life Technologies) and quantified via the Simple Neurite Tracing plugin in FIJI (NIH, U.S.) (Fanti et al., 2008). Specifically, the length of the longest bifurcation branch of each neuron was measured to characterize the neurite length of each neuron.

### 2.4 Immunostaining

To characterize the completion of differentiation, all differentiated cells were fixed with 4% paraformaldehyde (Thermo Fisher, U.S.) followed by permeabilization overnight in 1% Triton-X (Sigma, U.S.). Cells were then stained with anti-MAP2 antibody (1: 200 dilution, PA5-17646, Invitrogen) at 4 °C overnight followed by staining with a secondary goat-anti-rabbit antibody coupled with Alexa 568 (ab175471, Abcam, US).

Immunostaining for selected epigenetic modifications, namely 5mC, H3K9me3 and H3K27me3, was carried out following a similar protocol as described above. An additional denaturation step was included for 5mC staining, in which the cells are treated with 4N HCl for 30 minutes and equilibrated with 100mM Tris-HCl (pH 8.5) for 10 minutes. 5mC (61479, Active Motif, U.S.), H3K9me3 (ab8898, Abcam, U.S.) and H3K27me3 (ab192985, Abcam, U.S.) antibodies were used as primary. Goat-anti-mouse coupled with Alexa 488 (R37120, Thermo Fisher, U.S.) and Goat-anti-rabbit coupled with Alexa 568 (ab175471, Abcam, US) were used as secondary antibodies for 5mC and histone modifications, respectively.

### 2.5 Fluorescent microscopy

All fluorescent images were collected using a Nikon Eclipse Ti-2 inverted microscope. A 60×/1.40 NA oil objective was used for all images. Z-stack of cells were collected using Nikon EZ-C1 software with an interval of 1 μm. 2D max projections were created with ImageJ as we described previously(Sánchez et al., 2019; Sánchez et al., 2020). All images were then analyzed using a customized Cell Profiler pipeline similar as we described in our previous publication(Sánchez et al., 2020).

### 2.6 Gene expression analysis via RT-qPCR

Pellets of SH-SY5Y cells were collected immediately after completion of Pb treatments (96 hrs.) to determine changes in key epigenetic enzymes, namely DNMT1, DNMT3A, DNMT3B and TET1 for 5mC; KMT1A and KDM4A for H3K9me3, and EZH2, KDM6A and KDM6B for H3K27me3. β-Actin (ACTB) was used as the reference gene. Total RNA was extracted using an RNA purification kit (PureLink, Thermo Fisher Scientific, U.S.) following the manufacturer’s protocol and then reverse transcribed using SuperScript IV Reverse Transcriptase (Invitrogen, U.S.) with random hexamer primers. PowerUp SYBR Green (Applied Biosystems, U.S.) was used. Quantitative PCR (qPCR) was performed using QuantStudio 3 (Thermo Fisher, U.S.) following the MIQE guideline(Bustin et al., 2009). All qPCR primers were summarized in **Table S2** (Supporting Information).

### 2.7 Data analysis and statistics

All data were presented as mean ± standard deviation unless otherwise specified with independent replicates ≥ 3. All statistical analysis and calculations were performed using OriginPro (Version 2019, North Hampton, MA) statistical software. Analysis of variance and Tukey’s post-hoc test was used to calculate significance at different *p*-values. Principle Component Analysis and *k*-means clustering analysis was performed using Jupyter Notebook.

## 3. RESULTS

### 3.1 Exposure to low dose of Pb affects SH-SY5Y differentiation

SH-SY5Y cells were treated with 0, 15 and 50 ppb of Pb prior to differentiation for 96 hours (4 days) following the scheme outlined in **Fig. 1A**. The dose of Pb was selected based on the current regulation standard while accounting for Pb concentrations observed in recent public health incidences. Specifically, the maximum allowed level of Pb in water is 15 ppb as set by EPA. Higher Pb concentrations have been recently reported in several public health incidence related to Pb exposure(Triantafyllidou et al., 2009; Triantafyllidou et al., 2013) Furthermore, U.S. Center for Disease Control and Prevention has recommended BLL of 5μg/dL (50 ppb) in children as the action level that requires medical attention.

We started by examining the effects of selected Pb concentrations on cell viability and morphology. After 96 hours of treatment, the selected Pb doses were found to have minimal effects on cell viability (**Fig. 1B**) and morphology, including nuclear size, eccentricity (which measures ratio of major and minor axis, e=0 is a circle) and extent (which quantifies the irregularity of the nucleus, protrusions results in larger extents) (**Figs. S1A-C**, (Supporting Information)) consistent with previous literature reports suggesting low doses of Pb (< 1 μM ≅ 207 ppb) have minimal effects on cell viability and phenotype (Crumpton et al., 2001; Deng et al., 2001).

Pb was then removed from culture medium by washing three times with PBS before starting differentiation. SH-SY5Y cells can be differentiated into neuron-like cells that stain positive for MAP2 (see **Figs. 1C** and **S2A** (Supporting Information)), a neuron-specific protein that is essential for neurogenesis (Teng et al., 2001; Chilton and Gordon-Weeks, 2007). In unexposed cells, ~ 75% of cells are found to be MAP2 positive (MAP2+%) (**Fig. S2B**, Supporting Information), which is in close accordance with literature reports suggesting the SH-SY5Y cells can be used as a model system to study neuronal differentiation (Dwane et al., 2013). Exposure to Pb can reduce MAP2+% particularly at 15 ppb as shown in **Fig. S2B** (Supporting Information). Although decrease in mean values are observed at 50 ppb compared to the control, the decrease was not found to be statistically significant. We subsequently quantified MAP2 expression as MAP2 staining intensity per cell. As shown in **Fig. 1D**, MAP2 intensity was significantly reduced after exposure to Pb. Quantitatively MAP2 intensity was reduced by ~ 30 and 10% after exposure to 15 and 50 ppb of Pb, respectively as shown in **Fig. 1D**.

Neurite outgrowth is an important characteristic of neuron that can be typically measured by neurite length and complexity. Neurite length was thus measured via ImageJ as shown **Fig. S2C** (Supporting Information) and summarized in **Fig. 1E**. Exposure to Pb has resulted in longer neurite length at both 15 and 50 ppb but with seemingly reduced level of complexity (see **Fig. S2D** (Supporting Information)).

Nuclear morphology, including nucleus area, eccentricity and extent of differentiated neurons were determined similarly as described previously as shown in **Figs. S1D-F** (Supporting Information). After the completion of differentiation, significant changes were observed in nucleus area and extent, suggesting the potential attenuation of exposure effects through differentiation.

### 3.2 Low-dose of Pb exposure results in significant acute changes in epigenome

Epigenetic changes consist primarily of DNA methylation and histone post-translational modifications. Compared to active epigenetic markers, i.e., histone acetylation, repressive epigenetic marks, such as DNA methylation (methylation of cytosine in particular), H3K9me3 and H3K27me3 are crucial for forming transcriptionally repressed regions essential for cell lineage determination(Hawkins et al., 2010; Becker et al., 2016) and tend to persist over time. We thus evaluated immediate changes in these repressive markers after Pb exposure.

SH-SY5Y cells were exposed to Pb for 96 hours, washed, fixed and immuno-stained for the selected epigenetic modifications, including 5mC, H3K9me3 and H3K27me3 as shown in **Figs. 2A-C**. More images of immuno-stained cells can be found in **Figs. S3A, D and G (Supporting Information)**. After exposure, undifferentiated SH-SY5Y cells stained with 5mC antibodies exhibit small islands enriched in 5mC as bright foci. Changes in 5mC can be quantified via <u>I</u>ntegrated <u>I</u>ntensity per <u>N</u>ucleus (IIN) as established in literature(Kageyama et al., 2007; Ramsawhook et al., 2017; Stefanovski et al., 2017). Upon visual examination, Pb treatment does not seem to immediately elicit significant changes in 5mC patterns. Analysis of IIN, however, suggests that Pb treatment can lead to ~ 12.3 and 8.5% reduction in 5mC levels for 15 and 50 ppb treated SH-SY5Y cells as shown in **Fig. 2D**.

**Fig. 2.**
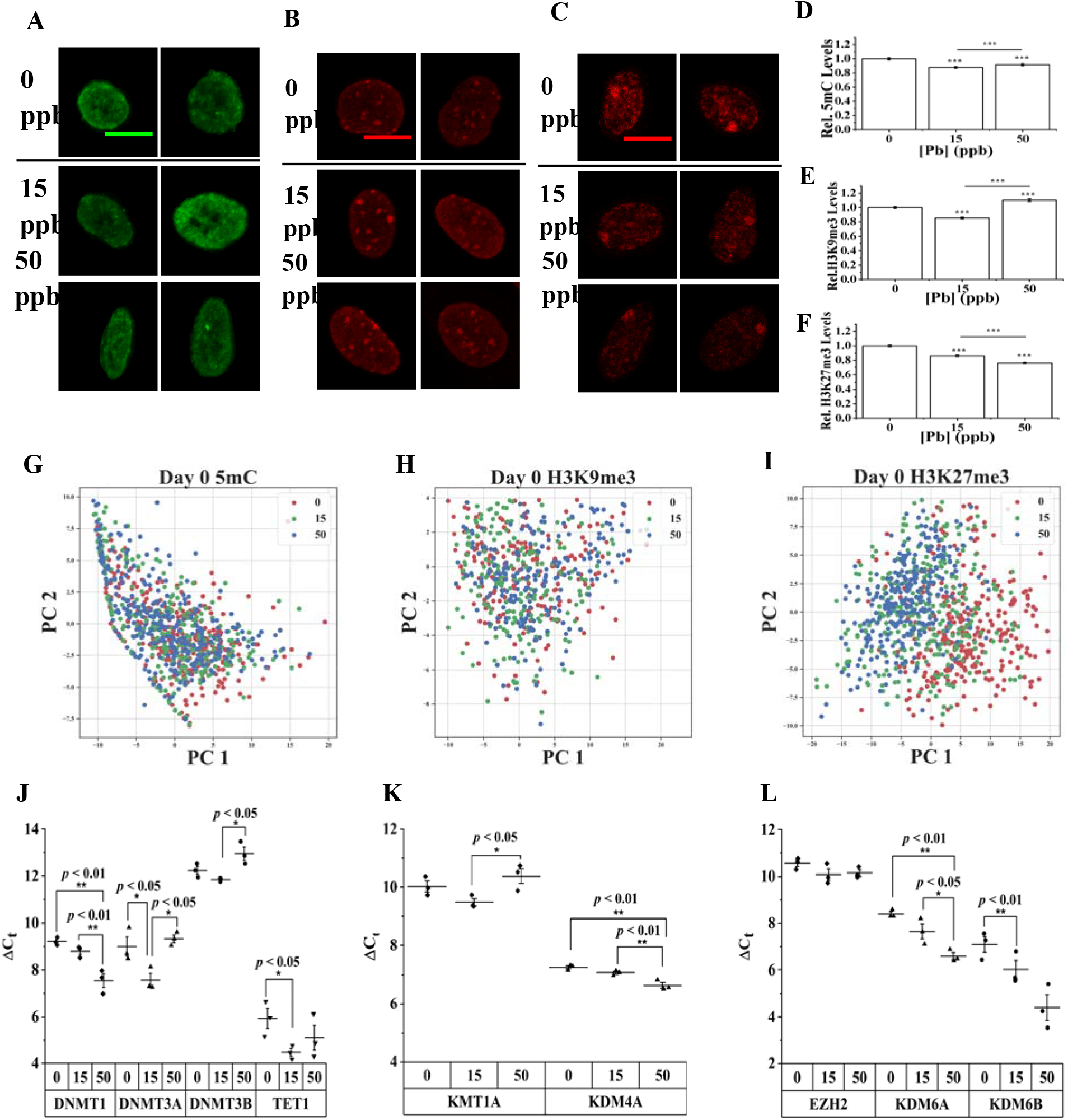
**(A-C)** 2D stacked confocal images of undifferentiated SH-SY5Y cells stained for **(A)** 5mC, **(B)** H3K9me3, and **(C)** H3K27me3 after 96 hrs exposure to Pb of varying doses. Scale bar = 10 μm. **(D-F)** Relative changes in epigenetic modifications, namely **(D)** 5mC, **(E)** H3K9me3 and **(F)** H3K27me3 after exposing to Pb of varying doses. Data = mean ± standard error. n > 500 cells. ***[=] p < 0.001. **(G-I)** Principal component analysis of texture features. Principal component 1 and 2 (PC1 and PC2, respectively) were selected as the axes explaining most of the data variance. **(G)** 5mC, **(H)** H3K9me3, H3K27me3. **(J-L)** Expression levels of mRNA coding for epigenetic writer and erasers, assessed immediately after 96 hours of exposure to Pb. Data are presented as the mean difference in cycle threshold (Ct) between gene of interest and β-actin. Error bars represent standard deviation. **(J).** Enzymes responsible for 5mC, namely DNMT1, DNMT3A, DNMT3B, and TET1. **(K).** Enzymes that are highly specific for H3K9me3, namely KMT1A and KDM4A. **(L).** Enzymes that are highly specific for H3K27me3, namely EZH2, KDM6A and KDM6B. Data = mean ± standard deviation. n > 3. * [=] p

A more in-depth pattern analysis was then carried out by identifying cell nucleus followed by determining foci features using a customized CellProfiler pipeline, as we detailed in our previous work(Sánchez et al., 2019). Representative images showing nucleus and foci identification via our pipeline for 5mC-stained images are shown in **Fig. S4A** and **B** (Supporting Information). To highlight changes in 5mC patterns due to Pb exposure, we included texture features only in our clustering analysis (all intensity and shape related features were omitted in constructing the principal component space). A PCA plot illustrating the separations of all treated groups are shown in **Fig. 2G**. No good separation is observed among differently treated cells suggesting no significant texture changes consistent with our visual observations.

RT-qPCR was then performed to determine changes in epigenetic enzymes that modulate 5mC levels in cells. **Table S3** (**Supporting Information**) summarizes our qPCR results. Relative changes in the mRNA levels of DNA methyltransferase, including DNMT1, DNMT3A, and DNMT3B, and DNA demethylase TET1 are shown in **Fig. 2J**. Pb treatments significantly upregulate the transcription level of DNMT3A (*de novo* DNA methyltransferase) and TET1 at 15 ppb, and DNMT1 (maintenance DNMT) at 50 ppb with respect to the control, having fold changes in the gene expression of DNMT3A, DNMT1 and TET1 of 2.79, 3.32, and 2.78, respectively. In addition, within Pb treatments the transcription level of DNMT1, DNMT3A, and DNMT3B (*de novo* DNA methyltransferase) presented significant difference. The increment of Pb concentration from 15 ppb to 50 ppb thus caused an upregulation in DNMT1, while DNMT3A and DNMT3B were downregulated. These results can partially explain the obtained changes in 5mC during Pb exposure.

Similar analysis was performed for H3K9me3 and H3K27me3 stained cells. Briefly, H3K9me3 form larger and more distinctive clusters compared to 5mC. These clusters are enriched within cell nucleus marking constitutive heterochromatin regions as well as in the perinuclear region marking laminin-associated domains as shown in **Fig. 2B** and **S3D** (Supporting Information). Intensity analysis suggests ~ 12.3% reduction and 10.4% increase in H3K9me3 levels for cells exposed to 15 and 50 ppb of Pb, respectively as shown in **Fig. 2E**. Visual inspection of stained cells does not suggest any significant pattern changes. PCA analysis was performed using identified cluster features (see also **Fig. S4C** (Supporting Information)) as shown in **Fig. 2H**. Cells with different treatments are not distinguishable within the PC space suggesting no significant pattern changes. **Fig. 2K** summarizes changes in H3K9me3 writer (KMT1A) and eraser (KDM4A). Relative to the untreated control, Pb treatment mainly affects the transcription level of KDM4A, increasing its gene expression level by 1.52-fold at 50 ppb. Within Pb treatment groups, increasing the Pb concentration from 15 to 50 ppb causes a significant reduction in the mRNA level of KMT1A while the mRNA level of KDM4A was increased. The observed changes in epigenetic enzymes cannot account for the observed changes in H3K9me3 resulting from 15 ppb treatment.

H3K27me3-stained cells exhibit a bright locus near the nucleus periphery, corresponding to bar body found in female origin cells, i.e., SH-SY5Y, representing inactive X chromosome (Chadwick and Willard, 2004; Shenoda et al., 2018). Small foci enriched in H3K27me3 features are also found primarily within cell nucleus but also in the periphery regions as shown in **Figs. 2C and S3G** (Supporting Information).Intensity analysis of exposed cells revealed ~ 13.9 and 23.8% reduction in H3K27me3 levels for 15 and 50 ppb treatments, respectively as shown in **Fig. 2F**. Texture features from the segmented nucleus and foci (**Fig. S4D)** were compiled and analyzed using PCA as shown in **Fig. 2I**. Visual inspection of the plot suggests potential separations of Pb-naïve and treated cells, but the distinctions are not sufficient to form distinctive clusters using K-means clustering approach. RT-qPCR was then performed to assess changes in H3K27me3 writer (i.e., EZH2) and erasers (i.e., KDM6A and KDM6B) as shown in **Fig. 2L**. Relative to the control, exposure to 50 ppb of Pb increased the transcription level of KDM6A and KDM6B by ~ 3.6 and 6.6 folds, respectively. Within Pb treatments, mRNA levels for KDM6A significantly increased from 15 to 50 ppb. Conversely, no statistical difference was observed between the control and Pb treated groups for mRNA levels of EZH2. The observed changed in the expression of epigenetic enzymes can thus at least partially account for the observed changes in H3K27me3 after Pb exposure.

### 3.3 Epigenome changes remain in exposed SH-SY5Y cells after cessation of exposure and completion of differentiation

Pb was removed from SH-SY5Y cell culture prior to initiating the differentiation protocol. Differentiation was completed on Day 14 and cells were fixed followed by immuno-staining for 5mC, H3K27me3 or H3K9me3 then. Typical images of post-differentiation neurons are shown in **Figs. 3A-C** and **S3B, E**, and **H** (Supporting Information). Significant intensity changes are still observed among different treatment groups and were compared to immediately after exposure to Pb (Day 0) as summarized in **Figs. 3D-F**. Briefly, after differentiation, changes in 5mC are further amplified with a decrease in 5mC levels of ~ 55.1 and 46.4% for 15 ppb and 50 ppb treatment groups respectively **(Fig. 3D)**. The difference, however, is less discernable within Pb-treated groups. PCA analysis was carried out using texture features only as shown in **Fig. 3G**. The texture features of 5mC enable separations between Pb-naïve and treated groups after the removal of Pb and completion of differentiation. Two major clusters can be identified via Silhouette plot (Rousseeuw, 1987) followed by k-means clustering (LLOYD, 1982). Cluster 1 contains primarily untreated cells (~73 %) while Cluster 2 consists mainly of Pb-treated SH-SY5Y cells (~ 87 %) as shown in **Fig. S5A** (Supporting Information). The contributions of texture features were rank ordered via a Lasso analysis (Tibshirani, 2011) . Among the top 5 contributors as summarized in **Fig. S5B** (Supporting Information), entropy features of 5mC within a nucleus and texture features of each 5mC island (foci) were ranked the highest, suggesting association of significant arrangement of 5mC loci within a nucleus.

**Fig. 3.**
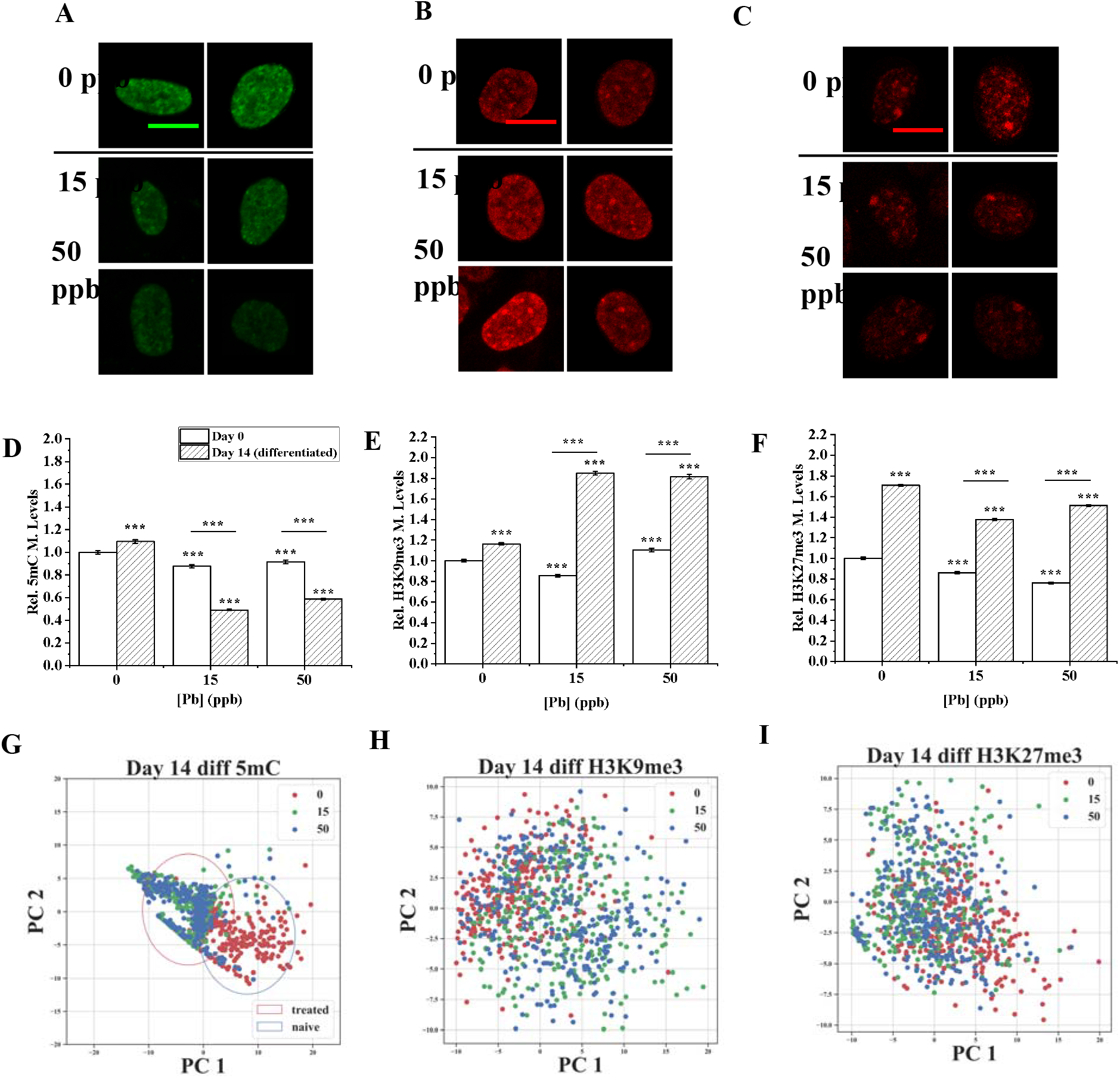
**(A-C)** 2D stacked confocal images of SH-SY5Y cells stained for **(A)** 5mC, **(B)** H3K9me3, and **(C)** H3K27me3 after 96 hrs exposure to Pb, removal of Pb by medium swapping and completion of differentiation. Scale bar = 10 μm. **(D-F)** Relative changes in epigenetic modifications, namely **(D)** 5mC, **(E)** H3K9me3 and **(F)** H3K27me3 after exposing to Pb of varying doses on Day 0 and Day 14 (after differentiation). All data are normalized to fluorescent intensity of untreated cells observed on Day 0. Data = mean ± standard error. n > 500 cells. ***[=] p < 0.001. **(G-I)** Principal component analysis of image pattern features. Principal component 1 and 2 (PC1 and PC2, respectively) were selected as the axes explaining most of the data variance. **(G)** 5mC, **(H)** H3K9me3, **(I)** H3K27me3.

Different from changes in 5mC, alterations in H3K9me3 and H3K27me3 are compensated and (partially) recovered after the completion of differentiation. Specifically, H3K9me3 level was increased by ~ 58.9 and 56.0%, respectively as compared to the untreated group in contrast to a global decrease right after exposure to Pb (**Fig. 3E**). Other than intensity, no significant texture changes are observed among treatment and control groups (**Fig. 3H**).

Differentiated SH-SY5Y cells exhibit ~19.4 and 11.4% decrease in H3K27me3 levels from pre-differentiation exposure to 15 and 50 ppb of Pb, respectively (**Fig. 3F**). The decrease in H3K27me3 levels was amplified for 15 ppb treated cells while a partial recovery was observed by cells exposed to 50 ppb contrasting to immediately after exposure (**Fig. 3F**). Texture analysis as shown in **Fig. 3I** does not suggest any significant alterations in H3K27me3 distributions within cell nucleus.

### 3.4 Differentiation as a potential confounding factor in propagating epigenetic memory

Systematic epigenome reprogramming is known to take place during differentiation leading to potential confounding effects (Li, 2002; Tibshirani, 2011). To understand this, control experiments were carried out as illustrated in **Fig. 4A**. Briefly, instead of starting differentiation right after exposure, SH-SY5Y cells were rinsed three times with PBS and continuously cultured in a Pb-free culture medium after cessation of exposure to Pb. Typical immuno-stained images for 5mC, H3K9me3 and H3K27me3 were summarized in **Figs. 4B-D** with more images in **Fig. S3 C, F** and **I** (Supporting Information). Differentiation results in significant increases in all selected repressive markers, namely 5mC, H3K9me3 and H3K27me3 levels as shown in **Fig. 4E-F** potentially as a result of heterochromatin establishment.

**Fig. 4.**
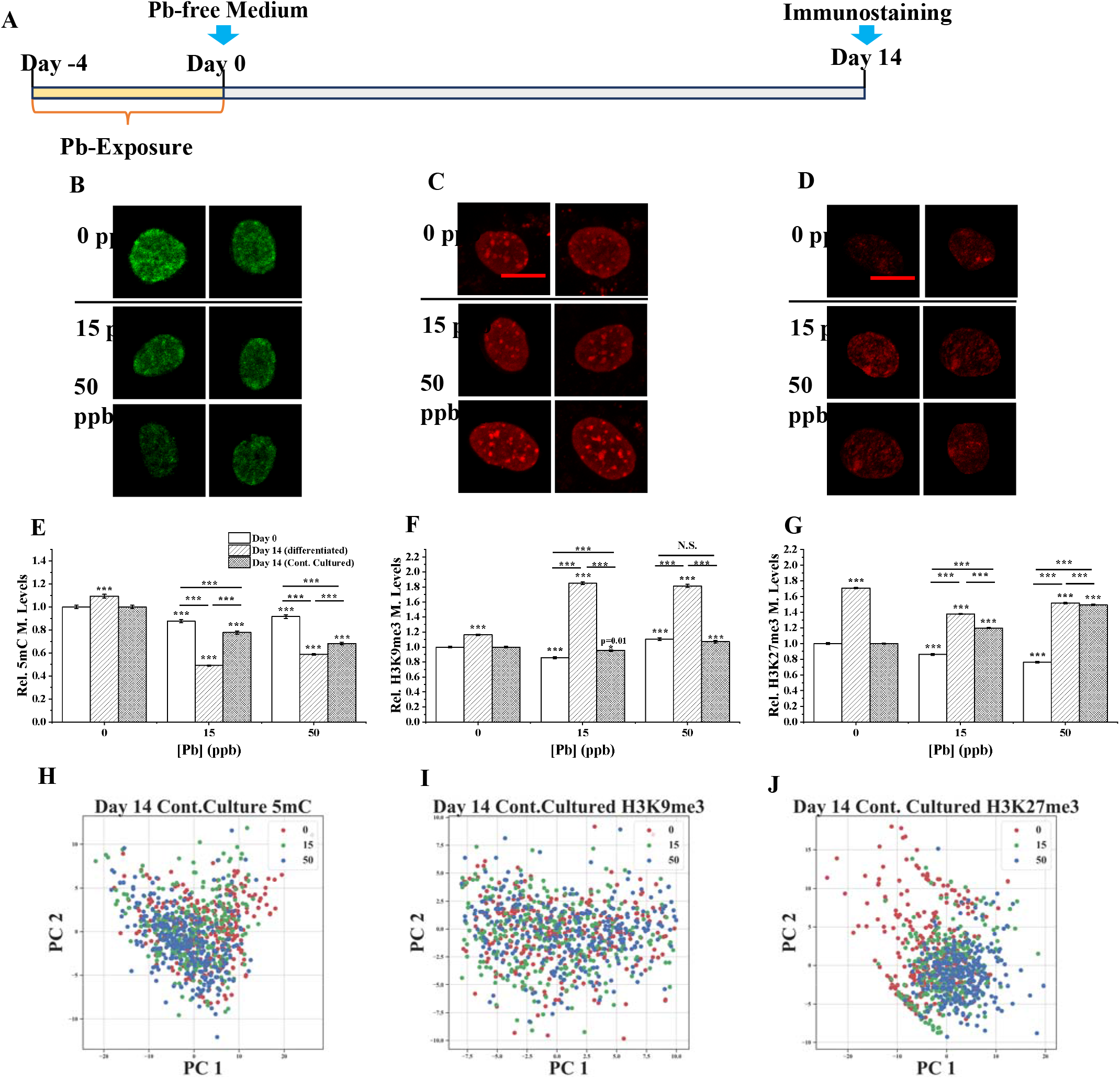
**(A)** A schematic illustration of the exposure and continuous culturing timeline for SH-SY5Y cells. (**B-D)** 2D projected immunostaining images of Pb exposed SH-SY5Y cells relaxed in Pb-free medium for 14 days **(B)** 5mC, **(C)** H3K9me3, **(D)** H3K27me3. Scale bar = 10 μm. **(E-F)** Bar graph demonstrating the relative changes in epigenetic modifications, **(E)** 5mC, **(F)** H3K9me3 and **(G)** H3K27me, after exposure to Pb^2+^ quantified at different time and conditions: day 0, day 14 with differentiation, and day 14 without differentiation. All data are normalized to fluorescent intensity of untreated cells observed on Day 0. Data are represented as mean ± standard error. n > 500 cells. Asterisks above the column indicate significance relative to the control, N.S.: no significance (p > 0.05), ***[=] p < 0.001, **(H-J)** Principal component analysis for image pattern features after continuous culturing. Principal component 1 and 2 (PC1 and PC2, respectively) were selected as the axes explaining most of the data variance. **(H)** 5mC, **(I)** H3K9me3, **(J)** H3K27me3.

After cessation of exposure to Pb for 14 days, other than 5mC, changes in H3K9me3 and H3K27me3 are both compensated for as summarized in **Figs. 4E-G**. Specifically, changes in 5mC after exposure to Pb are retained and amplified after 14 days. The magnitude in changes (~ 22.2 and 31.9% for 15 and 50 ppb of Pb, respectively), however, is much smaller compared to cells undergoing differentiation (~55.1 and 46.4% for 15 and 50 ppb of Pb, respectively) as shown in **Fig. 4E**. Changes in H3K9me3 are partially recovered after 14 days in continuously cultured SH-SY5Y cells (No statistical significance observed for 15 ppb, and ~7% increase for 50 ppb), while they were overcompensated in differentiated SH-SY5Y cells (see **Fig. 4F**). Surprisingly, in continuously cultured SH-SY5Y cells, changes in H3K27me3 are overcompensated (~ 19.7 and 49.4% for 15 and 50 ppb of Pb, respectively), which exhibit a trend opposite of differentiated cells. Texture analysis was carried out similarly as described in the previous sections with results summarized in **Figs. 4H-J**. While the 5mC, H3K9me3 H3K27me3 intensity levels of treated groups demonstrate significant difference from untreated controls, we cannot distinguish between Pb-naïve and treated based on their underlying texture features.

Changes in nuclear morphology of continuously cultured SH-SY5Y cells were also assessed as described for undifferentiated and differentiated cells previously. No significant changes were observed between Pb-naïve and treated groups as shown in **Figs. S1G-I** (Supporting Information).

## 4. DISCUSSION

### 4.1 Exposure to low dose of Pb elicits immediate changes in gene repressive markers

We selected to work with low doses of Pb (15 and 50 ppb) because of their relative high incidence of exposure (Hauptman et al., 2017; Lanphear et al., 2018) as well as increasing concerns of exposure-affiliated long-term neurodegenerative diseases at a later life stage (Shefa and Héroux, 2017; Bellinger et al., 2018). Exposure of SH-SY5Y cells to the selected doses of Pb for 96 hours do not elicit significant phenotypic changes in cells, including metabolic and morphological properties consistent with previous literature findings (Crumpton et al., 2001; Deng et al., 2001).

Meanwhile, significant changes are observed in selected epigenetic markers, namely 5mC, H3K9me3 and H3K27me3. 5mC is the most abundant epigenetic changes occurring on DNA and has an established role in gene regulation (Cedar and Razin, 2017; Kribelbauer et al., 2017). Exposure to Pb resulted in larger reductions in 5mC at 15 ppb compared to cells exposed to 50 ppb of Pb suggesting non-linear dependence in Pb doses. The acute 5mC changes occur globally with no significant pattern changes observed within cell nucleus right after exposure. Changes in the expression of 5mC writer and eraser enzymes are also observed with an overall increase in both writer and eraser enzyme expression at 15 ppb and dominate changes only in 5mC writer enzymes at 50 ppb. The expression level changes of these epigenetic enzymes can thus partially explain the observed 5mC but may also have contributions from altered epigenetic enzyme activities as demonstrated in literature (Sanchez et al., 2017).

Exposure to low dose of Pb has been shown to alter DNA methylation level in several animal studies, including zebrafish (Sanchez et al., 2017), rat (Sun et al., 2017; Singh et al., 2018a) and human cell lines (Li et al., 2011; Nye et al., 2015). The observed nonlinear dose-response is not uncommon. For example, in a recent rat study researchers have found that a significant larger number of genes are differentially methylated after exposure to low dose of Pb at 150 ppm as compared to 375 ppm (Singh et al., 2018b).

H3K9me3 and H3K27me3 are both suppressive markers typically found in transcriptionally silent chromatin regions. H3K9me3 are particularly enriched in constitutive heterochromatin that are almost permanently turned off (Becker et al., 2016). H3K27me3, on the other hand, is enriched in facultative heterochromatin that are suppressed and poised for potential activation(Richards and Elgin, 2002). Both H3K9me3 and H3K27me3 play an important role during differentiation(Tan et al., 2012; Tyssowski et al., 2014; Yao et al., 2016). For instance, protein-coding genes in an early stage, undifferentiated cells at germ-layer, exhibit high levels of H3K9me3 located mainly in gene bodies, promoters, and termination transcription sites that is reduced upon differentiation (Nicetto et al., 2019). The proper establishment of constitutive heterochromatin is relevant for preventing cell reprogramming and silencing undesirable lineage-specific genes to preserve cellular identity (Becker et al., 2016). H3K27me3 is more evenly distributed over genes and intragenic regions in differentiated cells and is reduced significantly during cell differentiation (Nicetto et al., 2019). Moreover, changes in H3K9me3 and H3K27me3 are known to be potentially persistent and inheritable across generations (Zenk et al., 2017; Perez and Lehner, 2019) warranting detailed study in this work. Global reduction in H3K9me3 level is observed at 15 ppb while a slight increase is observed at 50 ppb, mirroring discoveries in 5mC and suggesting a non-linear dose dependence. Although changes are observed in H3K9me3 writer and erase enzymes, the observed changes cannot fully explain the alterations in H3K9me3 levels at lower dosage Pb exposures. SH-SY5Y cells exposed to Pb exhibit significant reductions in H3K27me3 levels (with 13.9 and 23.8% for 15 and 50 ppb respectively). Significant increase is observed in the transcriptional level of H3K27me3 erasers, namely KDM6A and KDM6B providing a plausible explanation to the observed global changes in H3K27me3.

Limited studies exist examining the effects of Pb-exposure on histone modifications. A mouse study suggests that prenatal exposure to Pb of 100 ppm, where pups were continuous exposed to Pb from gestation to lactation, can reduce H3K9me3 level in the cortex of female and male rats by 49 and 65%, respectively (Schneider et al., 2016). Similar trend was recapitulated in our cell culture study. The observed changes in H3K9me3 seemingly mirror changes in 5mC at different Pb doses suggesting a potential cross-talking between DNA methylation and H3K9me3, which has been suggested in literature (Du et al., 2015; Zhao et al., 2016) and observed in our previous work of human cells exposure to atrazine (Sánchez et al., 2020).

Exposure to Pb (5μM ≅ 1035 ppb) has been reported to result in ~30% reduction in H3K27me3 at hippocampal neurons from rats (Gu et al., 2019). Similar reductions in H3K27me3 were also observed in hippocampus of female rats with chronic exposure to Pb of 125 ppm (Xiao et al., 2020). Our observations in SH-SY5Y cells are thus consistent with literature findings and thus can be plausibly explained by increase in the transcriptional level of H3K27me3 erasers.

### 4.2 Long-term epigenome changes with and without differentiation

Different from genetic mutations that are irreversible in nature, epigenetic changes are dynamic and thus potentially reversible. The dynamic balance of an epigenetic change is typically modulated by the presence of epigenetic writer and eraser enzyme. Most exposures to environmental chemicals are for a short duration of time. Assessing the persistence of acquired epigenome changes after cessation of exposure is thus critically important to understand their potential roles in long-term health. Here, we considered two different scenarios after cessation of exposure, namely through continuous culturing and via a neuron-specific differentiation. Changes persists in both cases while details vary depending on the type of modification and culturing methods.

After cessation of exposure, continuously cultured cells maintain its lower 5mC level compared to the untreated control and the changes are seemingly amplified after 14 days. Differentiation further amplifies the difference in 5mC levels among our experimental groups. In addition to global intensity changes, significant alterations in 5mC distribution patterns are also determined via texture analysis. After differentiation, untreated cells have significantly increased 5mC levels with more distinctive 5mC-enriched islands established within the cell nucleus. A more random and diffusive pattern, however, is observed in treated cells suggesting less well-defined heterochromatin regions separated by distinctive 5mC levels.

Different from 5mC, cells try to compensate for the loss in both H3K9me3 and H3K27me3 levels after removal of Pb with and without differentiation. H3K9me3 level is almost completely restored and becomes almost indistinguishable from the untreated group after 14 days of continuous culturing. With differentiation, significant increase in H3K9me3 is observed suggesting an overall upregulation in H3K9me3 likely prompted by homeostasis and differentiation driving forces. Changes in H3K27me3 are also compensated in both continuous culturing and differentiation experiments. The compensation, however, is more significant in continuously cultured cells resulting in an increase in H3K27me3 levels after 14 days, while differentiated SH-SY5Y cells with prior exposure to Pb becomes more like their untreated counterpart.

Accompanying the observed epigenome changes, we also noted significant alterations in nuclear size and morphology as summarized in **Fig. S1** (Supporting Information) only in differentiated cells treated with Pb. After exposure to Pb, differentiated neurons exhibit larger nucleus area relative to their untreated counterparts. Nuclear eccentricity that measures the roundness of cells were not significantly altered. Nuclear extent which quantifies shape irregularity was also found to be significantly altered after Pb exposure. The absence of nuclear alteration in after exposure and continuously cultured cells further suggests a potential compounding effect of low dosage Pb exposure on neuronal differentiation. Neurite length difference was also observed in differentiated SH-SY5Y cells after exposure to Pb with increased neurite length and reduce complexity. This finding suggests that neurite outgrowth was potentially impaired in differentiated neurons after exposure to Pb. This is consistent with previous study using PC12 cells suggesting Pb-treated PC12 cells can demonstrate enhanced neurite outgrowth (Crumpton et al., 2001; Davidovics and DiCicco-Bloom, 2005). Furthermore, exposed SH-SY5Y cells have lower neuron differentiation efficiency featuring decreased MAP2+% cells and the reduction in MAP2 expression levels suggests a less stable neuroarchitecture is formed. We then correlated changes in the observed phenotypic features (MAP2 expression and Neurite length) with Pb doses as shown in **Fig. 5**. Similar correlations were also performed using 5mC and Pb doses. A strong positive correlation was observed between the phenotypic alterations and 5mC changes (Pearson’s correlation = 0.9998 and 0.9891 for MAP2 expression and neurite length, respectively) suggesting a potential underlying correlative/causative relationship.

**Fig. 5.**
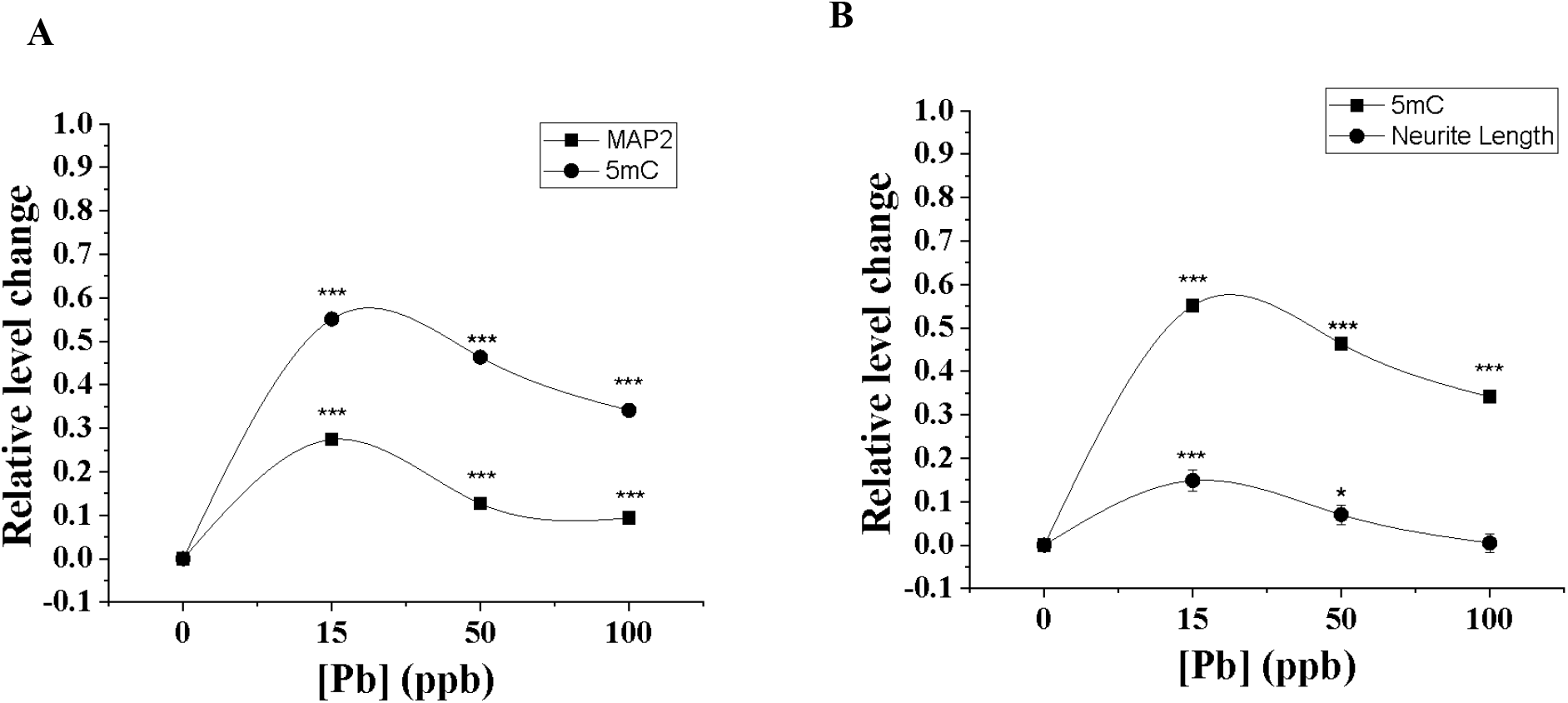
**(A)** Relative trends of changes in 5mC modification levels and MAP2 expressions on Day 14. Trends are positively correlated (Pearson’s correlation = 0.9998). **(B)** Relative trends of changes in 5mC modification levels and Neurite length on Day 14. Trends are positively correlated (Pearson’s correlation = 0.9891). An extra treatment concentration of 100ppb was added to enhance statistical power. 5mC modification levels, neurite length and MAP2 expression levels were quantified as described above. All data are normalized to untreated group. Data = Mean ± S.E. n > 500 cells. ***[=] p < 0.001.

Persistent changes in 5mC after exposure to Pb have been reported in PC12 cells using Pb doses of 50 (10.35 ppb), 250 (51.71 ppb), and 500 nM (103.5 ppb) (Li et al., 2012). Cynomolgus monkeys with Pb exposure at infancy was found to exhibit impaired DNMT activities (~20% reductions in activity) in brain tissue 23 years after exposure (Wu et al., 2008). Our observations are thus consistent with previous literature reports. To the best of our knowledge, no studies have evaluated the persistency of H3K9me3 and H3K27me3 marks after cessation of Pb exposure.

## 5. CONCLUSIONS

Collectively, exposure to low doses of Pb can alter 5mC levels which can persist and attenuate through time and differentiation. Changes in histone repressive markers, H3K9me3 and H3K27me3 are both compensated for during continuous culturing and differentiation. Over-compensations, however, are observed for H3K9me3 during differentiation and H3K27me3 during continuous culturing. Although no phenotypic changes are observed immediately after Pb exposure, significant alterations are observed in the differentiation ability, neurite outgrowth and neurite length after Pb exposure prior to differentiation. Dose-dependent phenotypic changes seem to correlate well with alterations in 5mC levels. Taken together, our results unequivocally suggest that exposure to low dose of Pb, particularly prior to differentiation, can result in persistent changes in epigenome last after cessation of Pb exposure and potentially contribute to the observed phenotypic changes with implications in long-term health.

## Notes

### Competing Interest Statement

The authors have declared no competing interest.

