## Supplemental files for "Low Dose Lead Exposure Induces Alterations on Heterochromatin Hallmarks Persisting Through SH-SY5Y Cell Differentiation"

**Supporting Tables:**

**Table S1.** Compositions of differentiation medium 1 and 2 and manufacturer

| Differentiation Medium 1 | |
| --- | --- |
| Components | **Provider** |
| EMEM | Gibco, U.S. |
| 2.5% hiFBS | Atlanta Biologicals, U.S. |
| 1% Pen/Strep | Thermofisher Scientific, U.S. |
| 10 µM Retinoic acid | Sigma Aldrich , U.S. |

| Differentiation Medium 2 | |
| --- | --- |
| Components | **Provider** |
| Neurobasal Medium | Gibco, U.S. |
| 1% B-27 | Thermofisher Scientific, U.S. |
| 1% Pen/Strep | Thermofisher Scientific, U.S. |
| 10 µM Retinoic acid | Sigma Aldrich, U.S. |
| 50 ng/mL BDNF | Sigma Aldrich, U.S. |
| 200 µM dibutyryl cyclic AMP | Sigma Aldrich, U.S. |

**Table S2.** Sequences of qPCR primers used in this work

| Gene Name | Primer Sequence 5’ – 3’ | Reference |
| --- | --- | --- |
| *DNMT1*  *DNMT3A*  *DNMT3B*  *TET1*  *KMT1A*  *KDM4A*  *EZH2*  *KDM6A*  *KDM6B*  *β - ACTIN* | Forward: GGTTTTCCTTCCTCAGCTACTGCGA  Reverse: CACTGATAGCCCATGCGGACCA  Forward: GGGGACGTCCGCAGCGTCACAC  Reverse: CAGGGTTGGACTCGAGAAATCGC  Forward: CCTGCTGAATTACTCACGCCCC  Reverse: GTCTGTGTAGTGCACAGGAAAGCC  Forward: CAGAACCTAAACCACCCGTG  Reverse: TGCTTCGTAGCGCCATTGTAA  Forward: GCACAAGTTTGCCTACAA  Reverse: CCAGGTCAAAGAGGTAGGTG  Forward: GAAGCCA CGAGCATCCTATGA  Reverse: GCGGAACTCTCGAACAGTCA  Forward: CCC TGA CCT TCT TAC TTG TGG  Reverse: ACG TCA GAT GGT GCC AGC AAT A  Forward: TACAGGCTCAGTTGTGTAACCT  Reverse: CTGCGGGAATTGGTAGGCTC  Forward: GGAGGCCACACGCTGCTAC  Reverse: GCCAGTATGAAAGTTCCAGAGCTG  Forward: GGAGTCCTGTGGCATCCACG  Reverse: CTAGAAGCATTTGCGGTGGA | [1]  [2]  [2]  [3]  [4]  [5]  [6]  [7]  [7]  [8] |

**Table S3.** The ΔΔC_t_, defined as the ΔC_t_ treated with ATZ compared to ΔC_t_ of untreated samples, with 95% confidence interval (CI) and expression fold change of selected genes in SH-SY5Y cells after ATZ exposure, 3 or 30 ppb, for 48 h. Statistical difference, *P* value (*P*), in the expression fold change against the control, SH-SY5Y cells Pb unexposed, was calculated using a one-way ANOVA followed by a Tukey’s HSD post-hoc test.

| Function of Epigenetic Enzymes | Target Gene | [Pb] ppb | ΔΔC_t_ ± SE | 95% CI | Fold Change | *P* |
| --- | --- | --- | --- | --- | --- | --- |
| 5mC writers | **DNMT1** | 15  50 | -0.41 ± 0.14  -1.67 ± 0.29 | -0.57 – -0.25  -2.00 – -1.34 | 1.35 ± 0.14  3.32 ± 0.70 | 0.826  0.017 |
|  | **DNMT3A** | 15  50 | -1.43 ± 0.28  0.33 ± 0.16 | -1.75 – -1.11  0.15 – 0.51 | 2.79 ± 0.48  0.80 ± 0.09 | 0.021  0.827 |
|  | **DNMT3B** | 15  50 | -0.39 ± 0.05  0.71 ± 0.27 | -0.45 – -0.33  0.40 – 1.02 | 1.31 ± 0.05  0.63 ± 0.11 | 0.159  0.078 |
| 5mC eraser | **TET1** | 15  50 | -1.45 ± 0.17  -0.81 ± 0.53 | -1.64 – -1.26  -1.41 – -0.21 | 2.78 ± 0.23  2.00 ± 0.63 | 0.048  0.268 |
| H3K27me3 writer | **EHZ2** | 15  50 | -0.47 ± 0.24  -0.40 ± 0.13 | -0.74 – -0.20  -0.55 – -0.25 | 1.43 ± 0.22  1.33 ± 0.12 | 0.204  0.355 |
| H3K27me3 erasers | **KDM6A** | 15  50 | -0.75 ± 0.31  -1.81 ± 0.14 | -1.10 – -0.40  -1.97 – -1.65 | 1.77 ± 0.38  3.55 ± 0.35 | 0.244  0.002 |
|  | **KDM6B** | 15  50 | -1.08 ± 0.39  -2.69 ± 0.54 | -1.52 – -0.64  -3.30 – -2.08 | 2.26 ± 0.51  6.62 ± 1.85 | 0.718  0.028 |
| H3K9me3 writer | **KMT1A** | 15  50 | -0.54 ± 0.12  0.36 ± 0.25 | -0.68 – -0.40  0.08 – 0.64 | 1.46 ± 0.12  0.81 ± 0.15 | 0.104  0.588 |
| H3K9me3 eraser | **KDM4A** | 15  50 | -0.18 ± 0.04  -0.60 ± 0.09 | -0.23 – -0.14  -0.70 – -0.50 | 1.13 ± 0.03  1.52 ± 0.10 | 0.380  0.003 |

**SUPPORTING FIGURES**

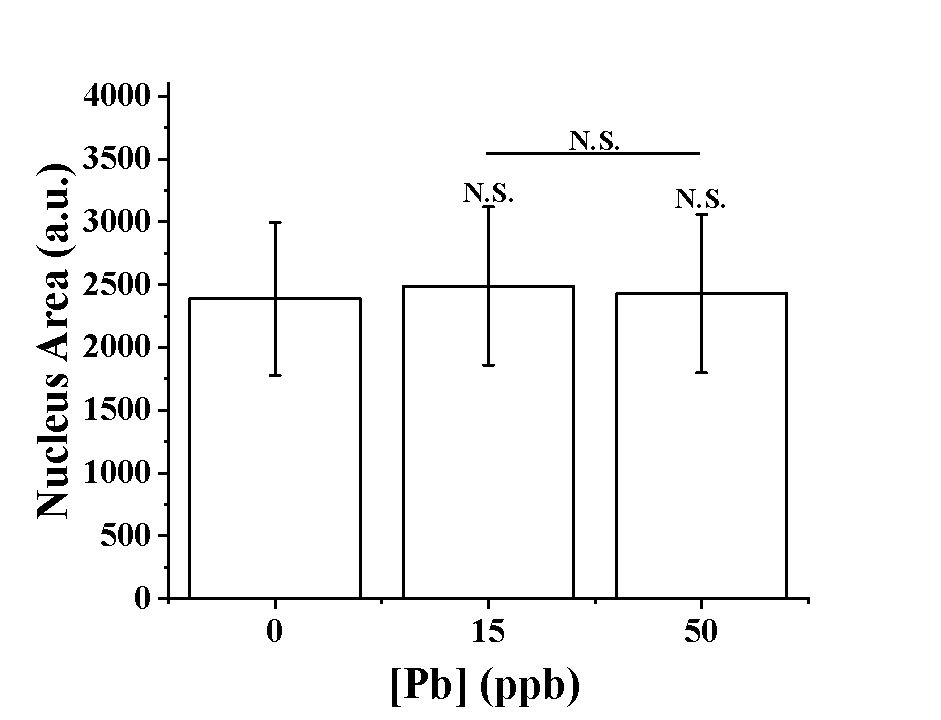

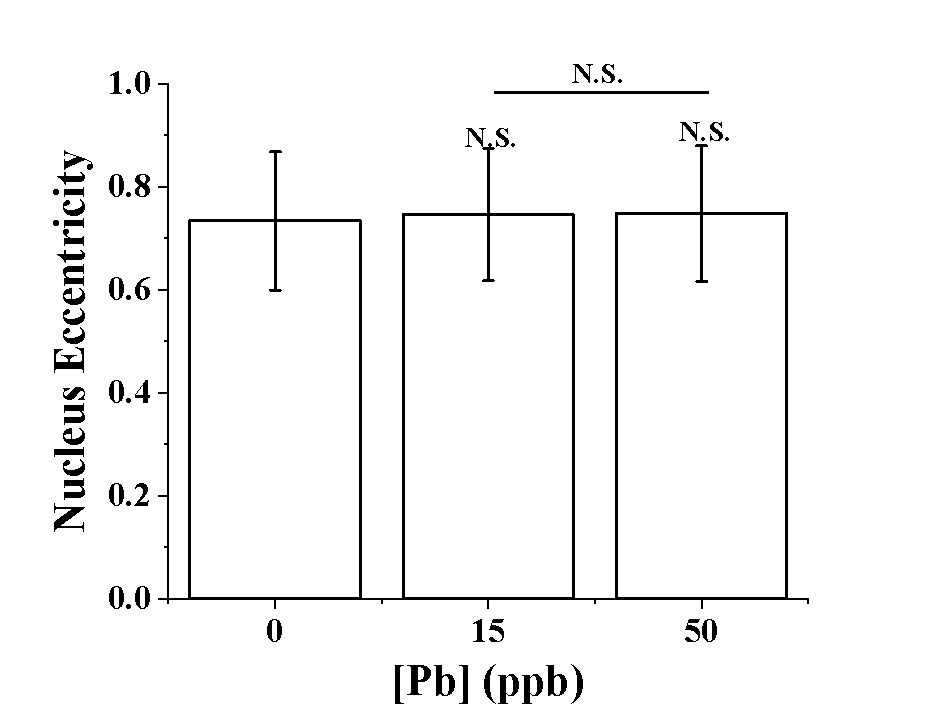

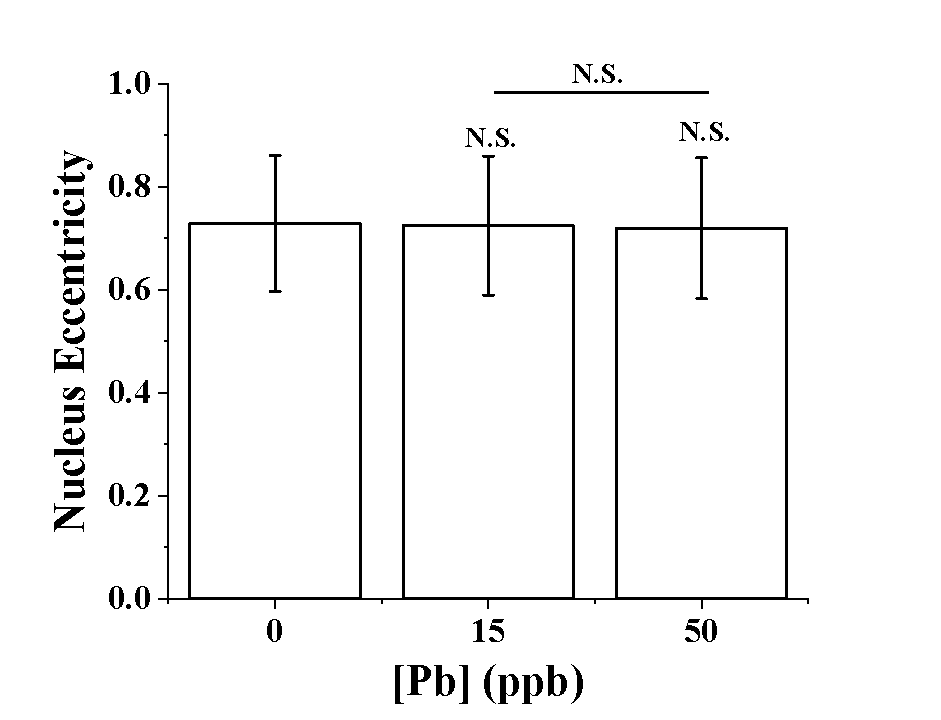

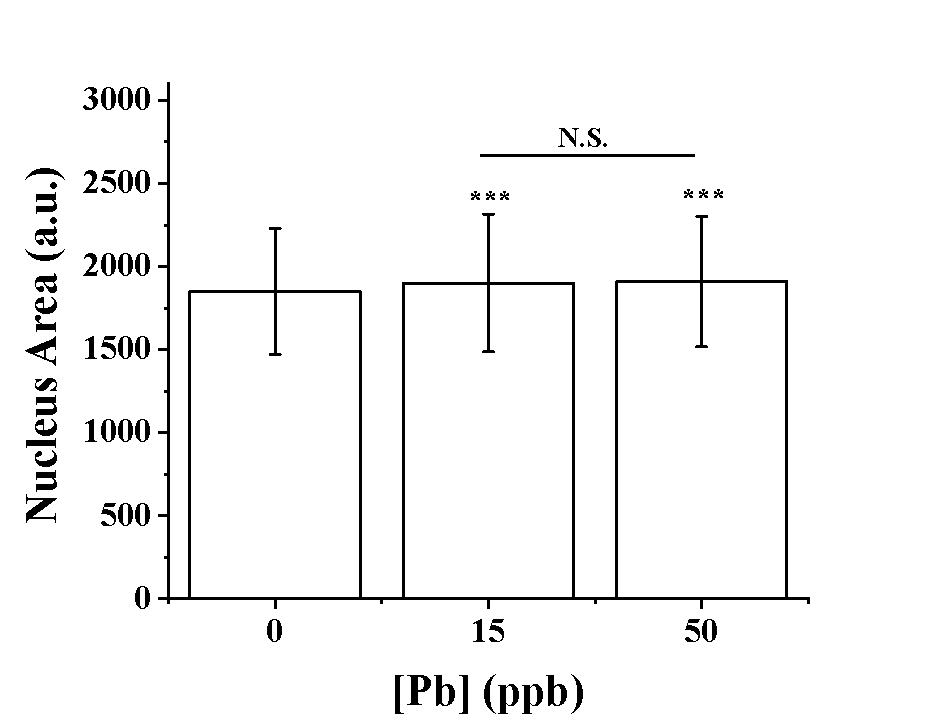

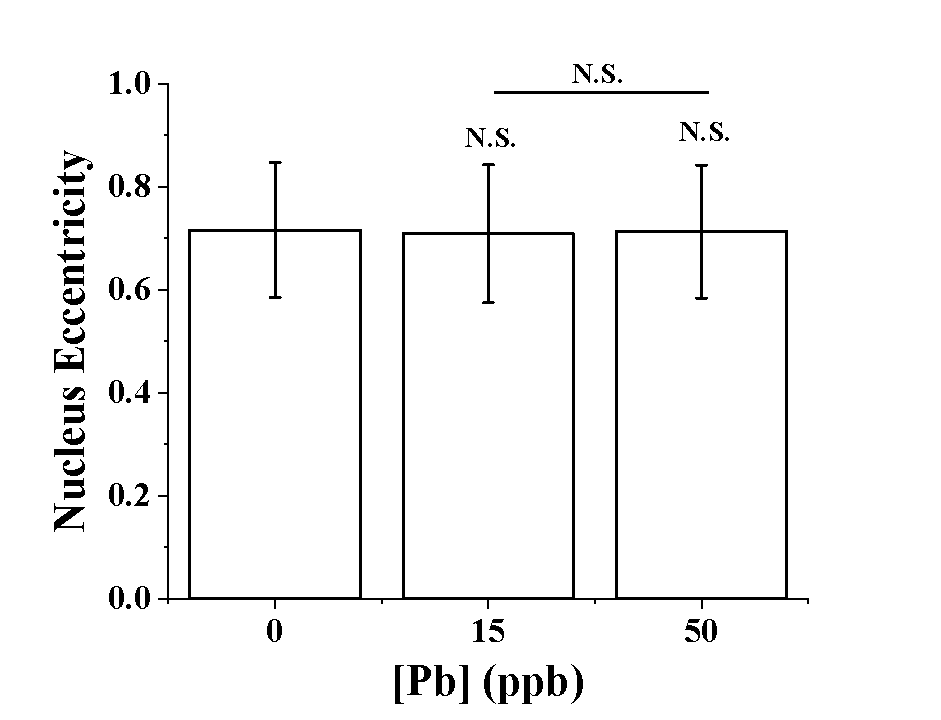

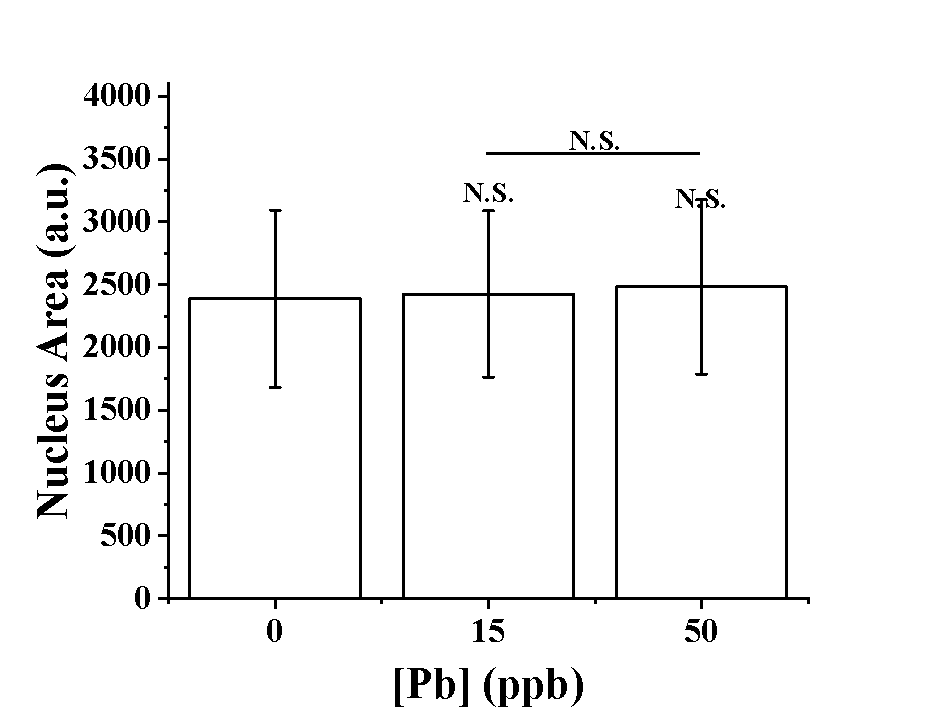

**A**

**B**

**Fig. S1**. Nuclear size **(A)**, eccentricity **(B)** and extent **(C)** of SH-SY5Y cells exposed to different concentrations of Pb for 96 hrs. Similar morphological information was collected for exposed cells after the completion of 14-day of differentiation **(D)-(F)** and continuous culturing **(G)-(I)**, respectively. n > 500 cells. Data = mean ± standard deviation.

**C**

**D**

**E**

**F**

**G**

**H**

**Day 0**

**Day 0**

**Day 14 differentiated**

**Day 14 differentiated**

**Day 14 Cont. Cultured**

**Day 0**

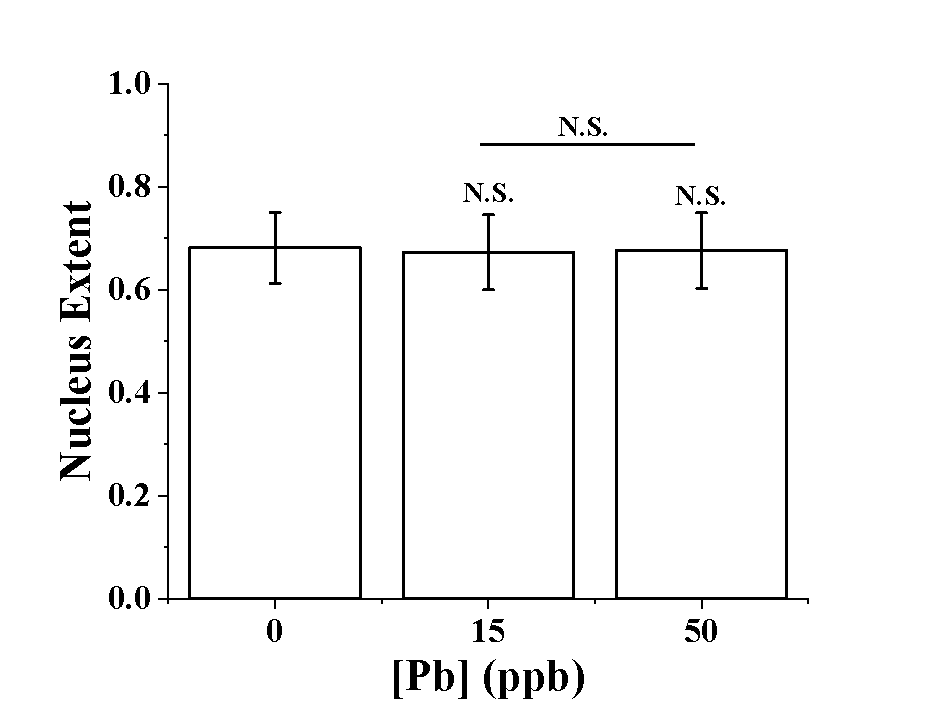

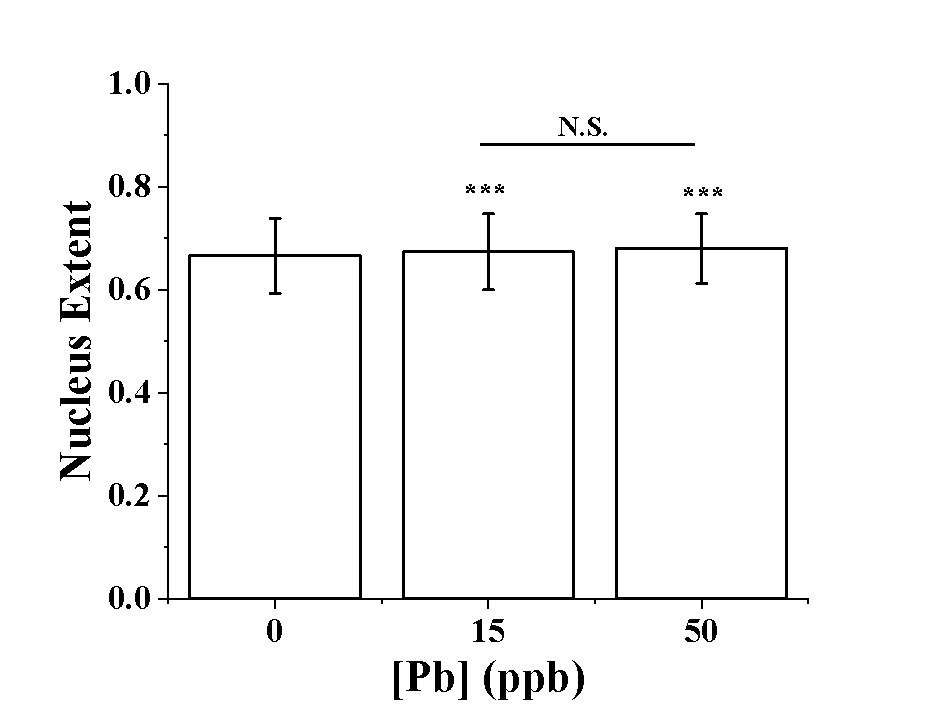

**Day 14 differentiated**

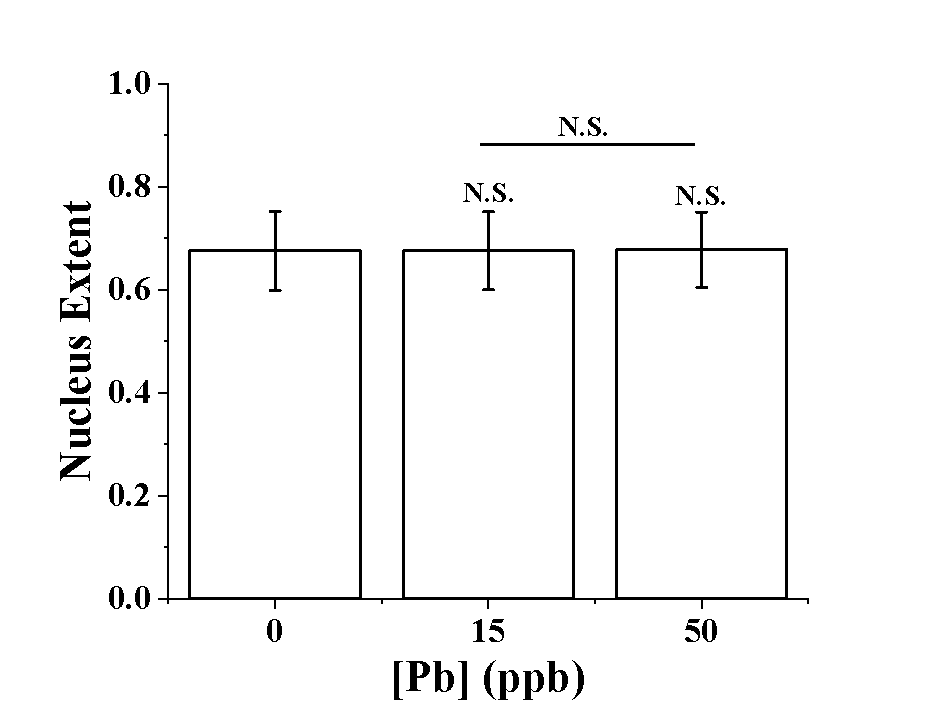

**Day 14 Cont. Cultured**

**Day 14 Cont. Cultured**

**I**

**Fig. S2**. **(A)** SH-SY5Y cells were exposed to Pb prior to the start of differentiation, fixed and stained for MAP2 (red) and nucleus (DPAI, blue). Scale bar = 50μm **(B)** MAP2+% of differentiated SH-SY5Y neurons after exposing to Pb. n = 3 **(C)** Typical neurite morphology determined in differentiated SH-SY5Y cells with Pb exposure prior to differentiation. Neurites are highlighted in red via Simple Neurite Tracing (FIJI).

**A**

**B**

**C**

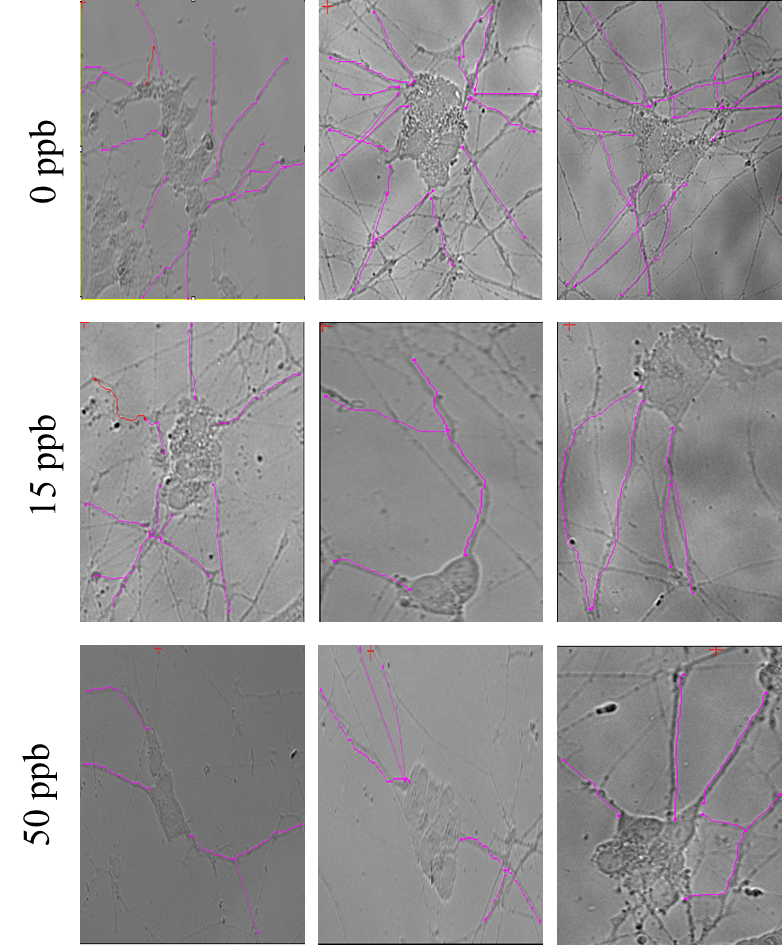

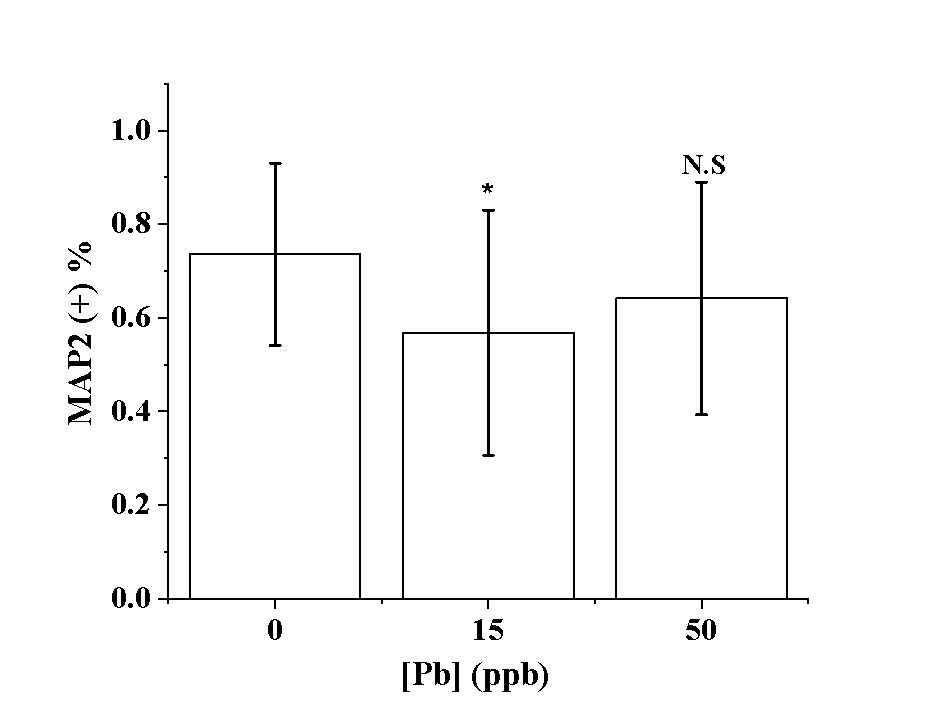

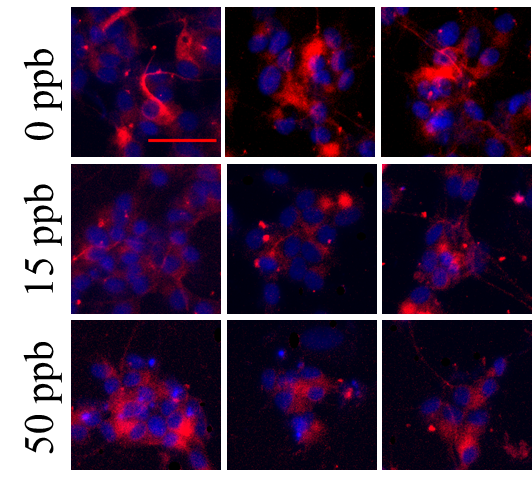

0 ppb

15 ppb

50 ppb

**A**

**B**

**Figure S3.** Representative of SH-SY5Y cells stained for selected epigenetic marks at Day 0 ((A), (D) and (G)), Day 14 after differentiation ((B), (E) and (H)) and Day 14 after continuous culturing( (C),(F)and (I)).Scale bar = 10 μm. **A, B, C**: 5mC; **D, E,F**: H3K9me3; and **G,H,I**: H3K27me3. All images are 2D confocal stacks via. max intensity projections.

**C**

**D**

**E**

**5mC**

**H3K9me3**

**H3K27me3**

**F**

**G**

**H**

**I**

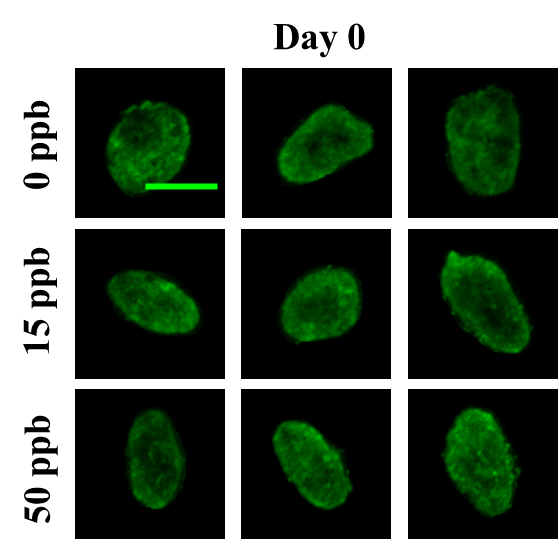

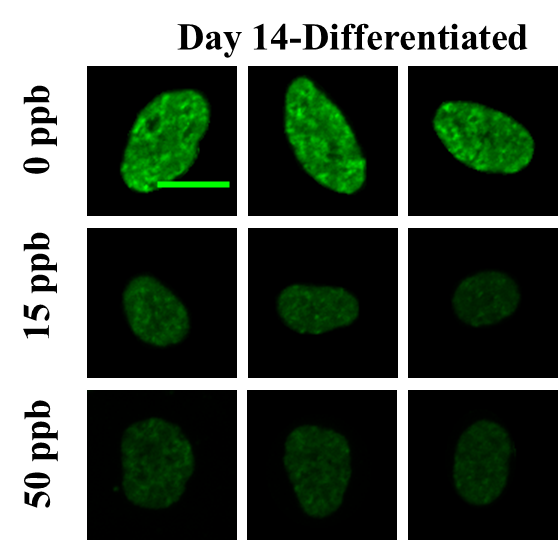

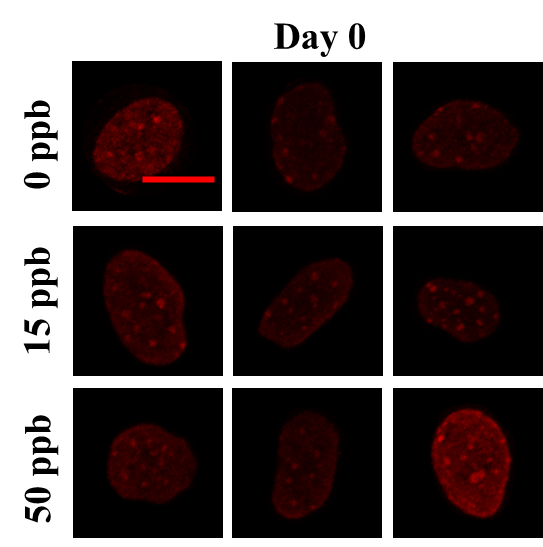

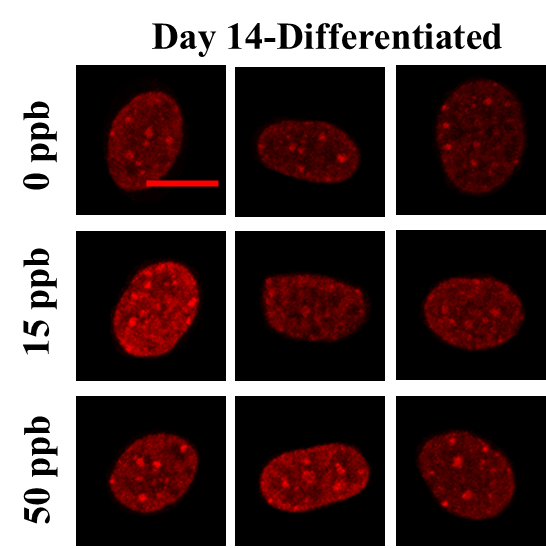

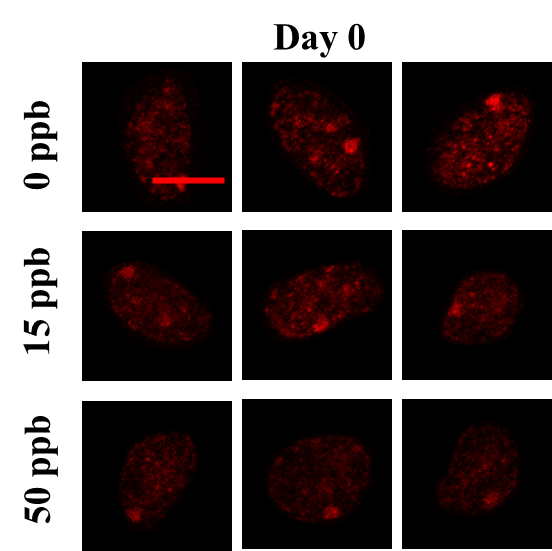

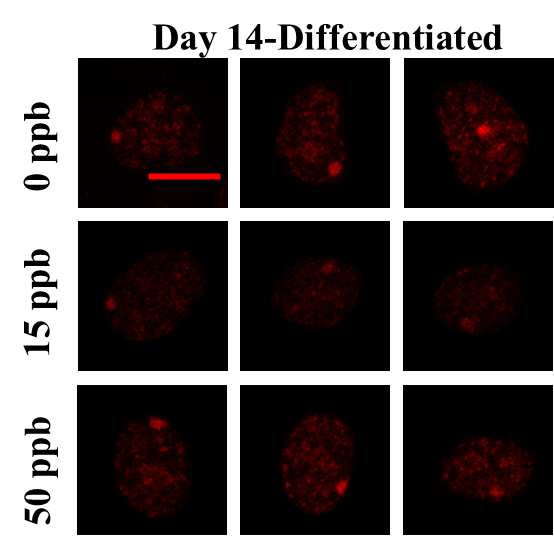

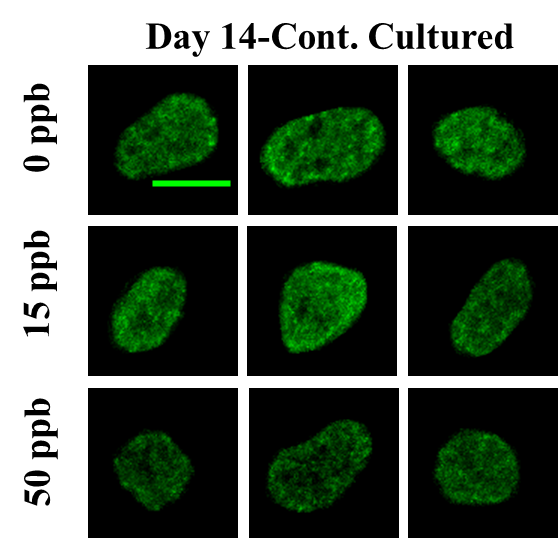

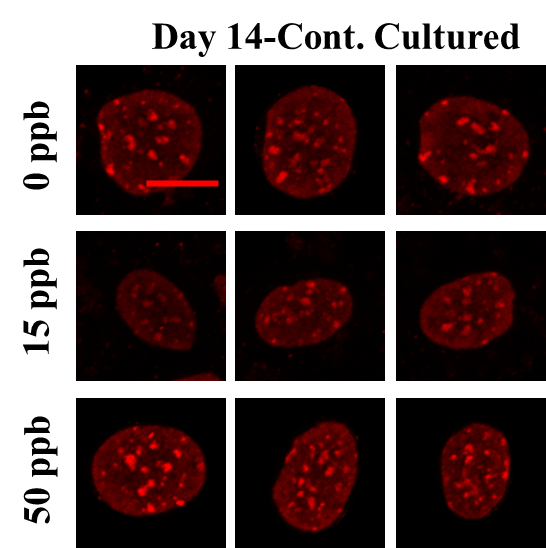

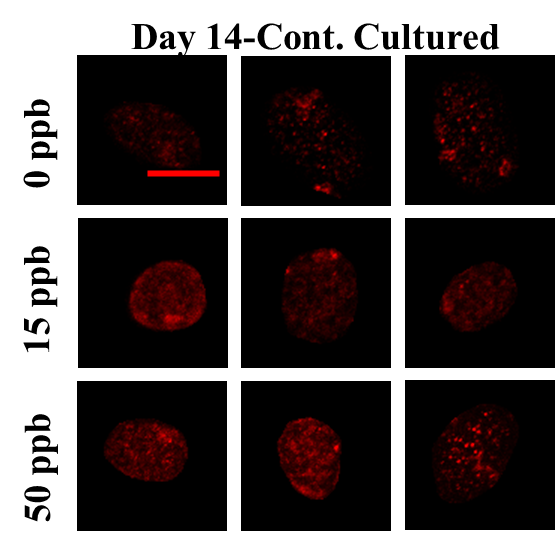

**Figure S4.** **(A)** Nuclei segmentation using Draq 5 staining. **(B-D)** Foci identifications of selected epigenetic markers using customized CellProfiler pipeline, **(B)** 5mC, **(C)** H3K9me3, and **(D)** H3K27me3.

**A**

**B**

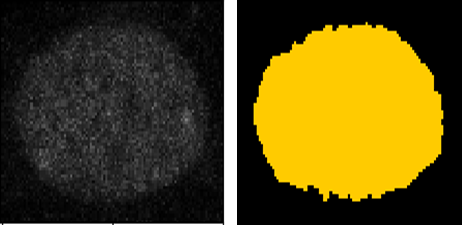

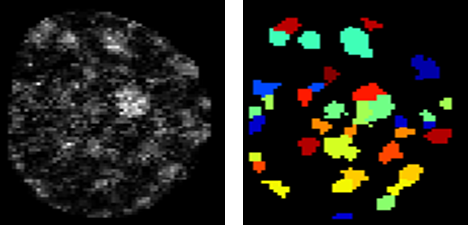

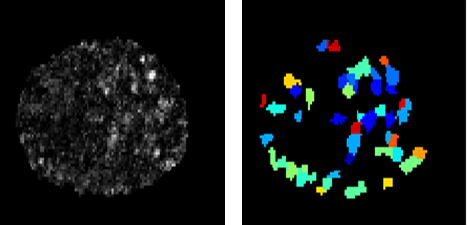

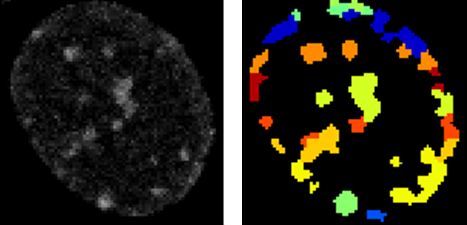

**C**

**D**

**Figure S5.** Percentage of differently treated cells within each cluster, **(A)** 5mC staining of differentiated neurons. **(B)** Rank ordered texture features that contribute to distinguishing between different clusters.

**A**

**B**

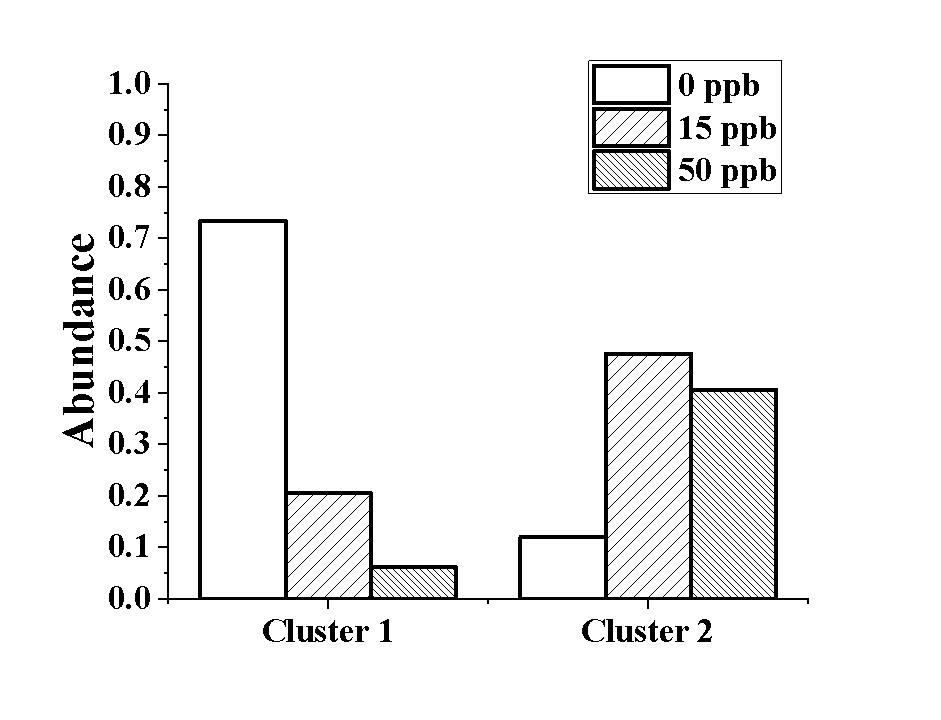

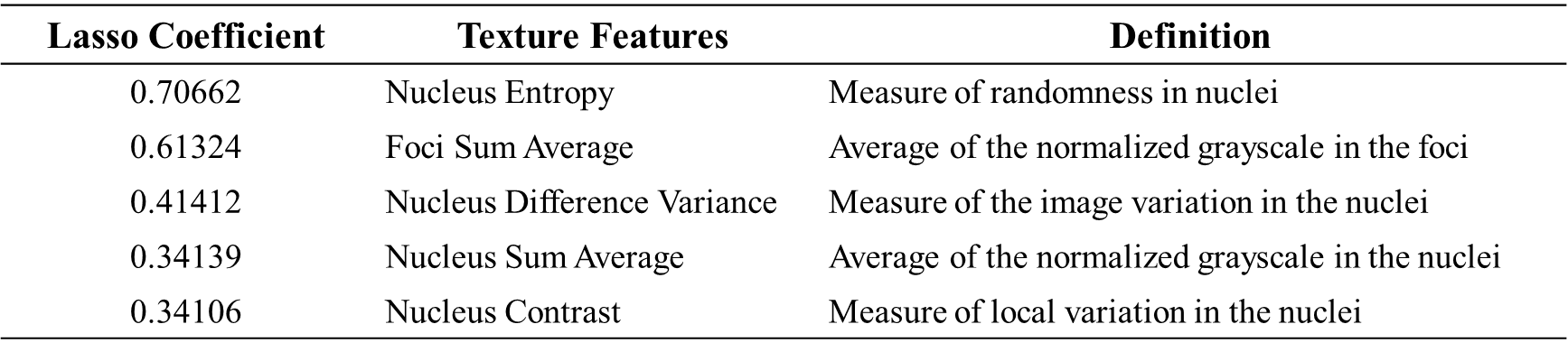
